# Dietary vs non-dietary fatty acid profiles of lake trout ecotypes from Lake Superior and Great Bear Lake: Are fish really what they eat?

**DOI:** 10.1101/714352

**Authors:** L Chavarie, J. Hoffmann, A.M. Muir, C.C. Krueger, C.R. Bronte, K.L. Howland, C.P. Gallagher, S.P. Sitar, M.J. Hansen, M.R. Vinson, L.F. Baker, L.L. Loseto, W. Tonn, H. Swanson

**Affiliations:** University of British Columbia, Biodiversity Center, 2212 Main Mall, Vancouver, BC, Canada, V6T1Z4; Scottish Centre for Ecology and the Natural Environment, Institute of Biodiversity, Animal Health & Comparative Medicine, University of Glasgow, Glasgow, UK; University of Waterloo, Department of Biology, 200 University Avenue West, Waterloo, Ontario, Canada, N2L 3G; Great Lakes Fishery Commission, 2100 Commonwealth Blvd., Suite 100, Ann Arbor, Michigan, USA, 48105; Michigan State University, Center for Systems Integration and Sustainability, 1405 South Harrison Road, 115 Manly Miles Building, East Lansing, Michigan USA 48823; U.S. Fish and Wildlife Service, Green Bay Fish and Wildlife Conservation Office, New Franken, WI, USA; Fisheries and Oceans Canada, 501 University Crescent, Winnipeg, Canada; University of Alberta, Department of Biological Sciences, CW 405 BioSciences Building, Edmonton, Canada; Michigan Department of Natural Resources, Marquette, MI, USA; U.S. Geological Survey Great Lakes Science Center, Hammond Bay Biological Station, USA (retired); U.S. Geological Survey Great Lakes Science Center, Ashland, WI, USA

**Keywords:** Intraspecific diversity, salmonid, aquatic ecology, trophic ecology, Great Lakes, *Salvelinus namaycush*

## Abstract

Fatty acids are well-established biomarkers used to characterize trophic ecology, food-web linkages, and the ecological niche of many different taxa. Most often, fatty acids that are examined include only those previously identified as “dietary” or “extended dietary” biomarkers. Fatty acids considered as non-dietary biomarkers, however, represent numerous fatty acids that can be extracted. Some studies may include non-dietary fatty acids (i.e., combined with dietary fatty acids), but do not specifically assess them, whereas in other studies, these data are discarded. In this study, we explored whether non-dietary biomarkers fatty acids can provide worthwhile information by assessing their ability to discriminate intraspecific diversity within and between lakes. Non-dietary fatty acids used as biomarkers delineated variation among regions, among locations within a lake, and among ecotypes within a species. Physiological differences that arise from differences in energy processing can be adaptive and linked to habitat use by a species’ ecotypes, and likely explains why non-dietary fatty acids biomarkers can be a relevant tool to delineate intraspecific diversity. Little is known about the non-dietary-mediated differences in fatty acid composition, but our results showed that non-dietary fatty acids biomarkers can be useful tool in identifying variation.

## Introduction

Constraints on traditional methods to investigate diets of organisms within aquatic systems have led to the development and use of biochemical tracers (Vinson and Budy 2010). Among these, fatty acids have gained popularity as both qualitative and quantitative trophic markers that reflect foraging patterns and food-web dynamics (Galloway et al. 2014, Iverson 2009). As such, fatty acids have been used to characterize variation within populations and species (e.g., evolutionary units linked to trophic polymorphism; Logan et al. 2000, Scharnweber et al. 2016), and to explore geographical (Hiltunen et al. 2016, Pomerleau et al. 2014, Quérouil et al. 2013) and temporal variation in trophic ecology (e.g., seasonal and annual variations; Eloranta et al. 2013, Hartwich et al. 2012).

Fatty acids are the “building blocks” of lipids and represent the largest constituent of neutral lipids (e.g., triacylglycerols and wax esters) and polar phospholipids (Iverson 2009). The array of fatty acids present in nature is exceptionally complex, routinely ~70 fatty acids can be identified within an organism (Budge et al. 2006, Iverson 2009). The utility of fatty acid analyses to reflect foraging patterns and food web dynamics relies on the assumption that lipids are broken down into their constituent fatty acids and incorporated relatively unchanged into consumer tissues (Howell et al. 2003, Iverson 2009, Iverson et al. 2004). The storage patterns of fatty acids depend on the biochemical limitations of organisms to biosynthesize, modify chain-length, and introduce double bonds into fatty acids, which culminate in vertebrates (Iverson 2009). Fatty acids that are stored in predator tissues, with no or little modification from their prey, have been labelled as “dietary” or “extended dietary” tracers (Budge et al. 2006, Iverson et al. 1997, 2004) and have been the target of analyses in ecological studies. Accordingly, for purposes of this study, we use the terminology of “dietary” versus “non-dietary” derived fatty acids based on the classification of Iverson et al. (2004), who identified typical dietary fatty acid markers that are now used across a variety taxa. Often, “non-dietary” fatty acids extracted from tissue samples are not examined, and only those recognized as dietary are analyzed. In other studies, dietary and non-dietary fatty acids may be combined to characterize food-web relationships, but information gained from inclusion of non-dietary fatty acids is not specifically addressed (Hiltunen et al. 2019, Mariash et al. 2017, McMeans et al. 2015, Taipale et al. 2019).

The exclusion or ignoring non-dietary fatty acids from analyses is not misguided, as the purpose of most studies is to describe diet patterns, and this practice has resulted in reliable information being produced across taxa (Galloway et al. 2014, Grosbois et al. 2017, Iverson 2009, Iverson et al. 2004). However, it is unknown whether valid information for other research questions is lost when investigators discard non-dietary fatty acids. Biological (e.g., phenotypic and genetic) and environmental variables can affect lipid and fatty acid composition in fishes (Olsen and Skjervold 1995), including temperature (Farkas et al. 1980, Olsen 1999), salinity (Borlongan and Benitez 1992), and light (Ota and Yamada 1971). Thus, fatty acids not labelled as “dietary” markers could be useful when the aim of a study is to delineate or better understand intraspecific diversity. To investigate this, we compared non-dietary to dietary fatty acids biomarkers among ecotypes of lake trout (*Salvelinus namaycush*) in Lake Superior and Great Bear Lake, as these ecotypes represent important intraspecific diversity in these lakes.

Salmonids, such as lake trout, inhabit young ecosystems believed to be 10,000 to 15,000 years old, e.g., post-glacial lakes colonized from non-glaciated refugia. The depauperate communities of post-glacial lakes are commonly characterized by reduced interspecific competition and predation, which allows colonizers access to a relatively wide array of resources, conditions that favour development of intraspecific diversity (McPhail 1993, Robinson and Wilson 1994, Smith and Skulason 1996). This high level of ecological opportunity, together with an increase in intraspecific competition after colonization, can promote specialization and divergence within a population, e.g., the development of groups of individuals with similar patterns of resource use (Svanbäck et al. 2007). Phenotypic characteristics of individuals in such groups may evolve as niches incorporate novel foraging resources, referred to as resource divergence (Robinson and Parsons 2002, Skulason and Smith 1995). As niches diverge, ecotypes can develop differences in morpholology, genetics, physiology, life-history, and/or behaviour (Bolnick et al. 2007, Schluter and McPhail 1992, Smith and Skulason 1996).

Intraspecific diversity in lake trout has been mostly linked to differences in depth distribution and, not surprisingly, is best known from large (> 500km^2^), deep lakes, such as Lake Superior (Moore and Bronte 2001, Muir et al. 2014), Lake Mistassini (Zimmerman et al. 2007), Great Slave Lake (Zimmerman et al. 2006), and Great Bear Lake (Chavarie et al. 2013). Although lake trout diversification has often focused on isolation-by-depth (without excluding isolation-by-adaptation) in large, deep lakes (e.g., Lake Superior), diversification also occurs in small lakes or within shallow-water habitats (Bernatchez et al. 2016, Chavarie et al. 2013, Chavarie et al. 2016c, Morissette et al. 2018).

A number of studies have assessed lake trout diets with fatty acids, either qualitatively (Chavarie et al. 2016b, Happel et al. 2017a, Happel et al. 2017b, Hoffmann 2017) or quantitatively (Happel et al. 2016a, Happel et al. 2016b), generating reliable information about dietary patterns. However, pronounced and systematic non-dietary-mediated differences in composition of fatty acids have been reported in lake trout, supporting the rationale that non-dietary fatty acids biomarkers could be important in delineating intraspecific diversity within this species. Goetz et al. (2013) found physiological differences between lean and siscowet ecotypes of lake trout that reflected genetically based differences in lipid synthesis, metabolism, and transport (Goetz et al. 2010), which suggests information from non-dietary fatty acids biomarkers might be important, especially if they have genetic-based mechanisms. Consequently, where intraspecific diversity is manifested as physiological differences among ecotypes, non-dietary fatty acids biomarkers could assist in identifying variation. Yet, little is known about the non-dietary-mediated differences in fatty acid composition of fish (or other taxa) in an ecological context.

To investigate the use of non-dietary fatty acids biomarkers as an ecological tool, we analyzed a set of non-dietary and dietary fatty acid biomarkers (as classified by the literature), from lake trout ecotypes collected from two lakes that sustain intraspecific diversity along different axes of intraspecific diversification (e.g., depth-dependent and depth-independent). Specifically, we compared: 1) non-dietary and dietary fatty acids biomarkers among four ecotypes of lake trout from each of Great Bear Lake and Lake Superior, and 2) how well non-dietary and dietary biomarkers delineated intraspecific diversity within and between lakes. Answers to these comparisons will help determine if non-dietary fatty acid biomarkers can discriminate fish geographically and among groups within a species, and if they offer different perspectives than dietary fatty acids biomarkers.

## Material and methods

### Study systems and intraspecific diversity of lake trout

Located in the northeast corner of Northwest Territories (Canada; 65°92′ N, 120°82′ W), Great Bear Lake is the most northerly lake of its size (~31 000 km^2^) and is the fifteenth deepest freshwater lake in the world (Fig. 1; Johnson 1975). A UNESCO biosphere reserve, Great Bear Lake is 250 km south of the Arctic Ocean and has characteristics typical of an Arctic lake. The lake is ultra-oligotrophic and, despite its size, has a simple food web, supporting only 15 fish species (Johnson 1975, MacDonald et al. 2004). The lake and its biota have remained relatively isolated and unexploited and is one of the most pristine large lakes in North America. Great Bear Lake has five semi-isolated arms, but due to sample sizes, data were pooled across multiple sites (see Chavarie et al. 2016b for details).

**Fig. 1.**
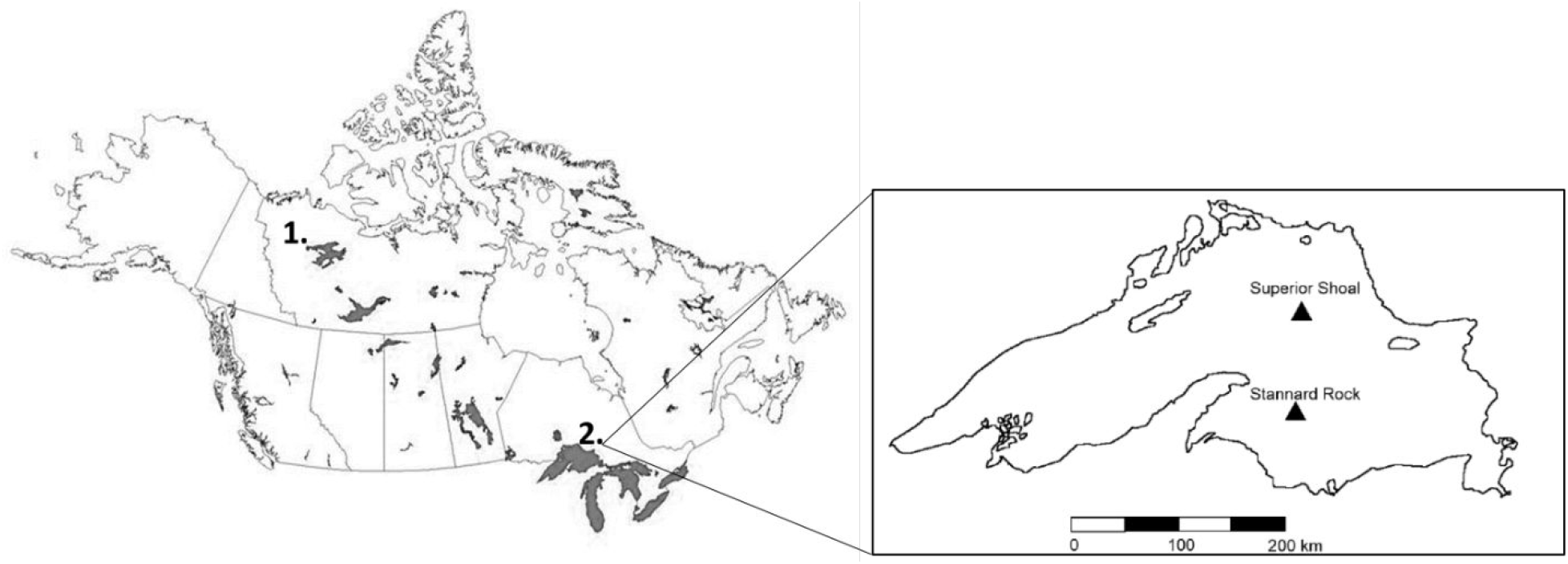
Location of (1) Great Bear Lake (Canada) and (2) Lake Superior (Canada-USA). Two sampling sites were defined in Lake Superior to account for spatial variation: Superior Shoal and Stannard Rock (QGIS 3.0, Canada Basemap; Hoffmann 2017).

Great Bear Lake sustains a noteworthy example of lake trout divergence (Fig. 2). With its intraspecific diversity independent of depth-based segregation, the lake also presents an unusual ecological framework for lake trout differentiation (Chavarie et al. 2016a). Currently, four shallow-water ecotypes are described in Great Bear Lake; three are common (Ecotypes 1-3), and one is rare (Ecotype 4). Ecotype 1 has the smallest head and jaws, intermediate fin lengths and body shape intermediate between Ecotypes 2 and 3. The lean-like Ecotype 2 has the largest head and jaws but the smallest fins, and a streamlined body shape. Ecotype 3 has the longest fins, a robust body shape, and a sub-terminal mouth. Ecotype 4 has a thick and curved lower jaw, a streamlined body shape, and the smallest caudal peduncle among the ecotypes. Although Ecotype 4 is a pelagic specialist, Ecotypes 1-3 have more general feeding habits, with varying degrees of omnivory along a weak benthic-pelagic gradient (Chavarie et al. 2016a, Chavarie et al. 2016b).

**Fig 2.**
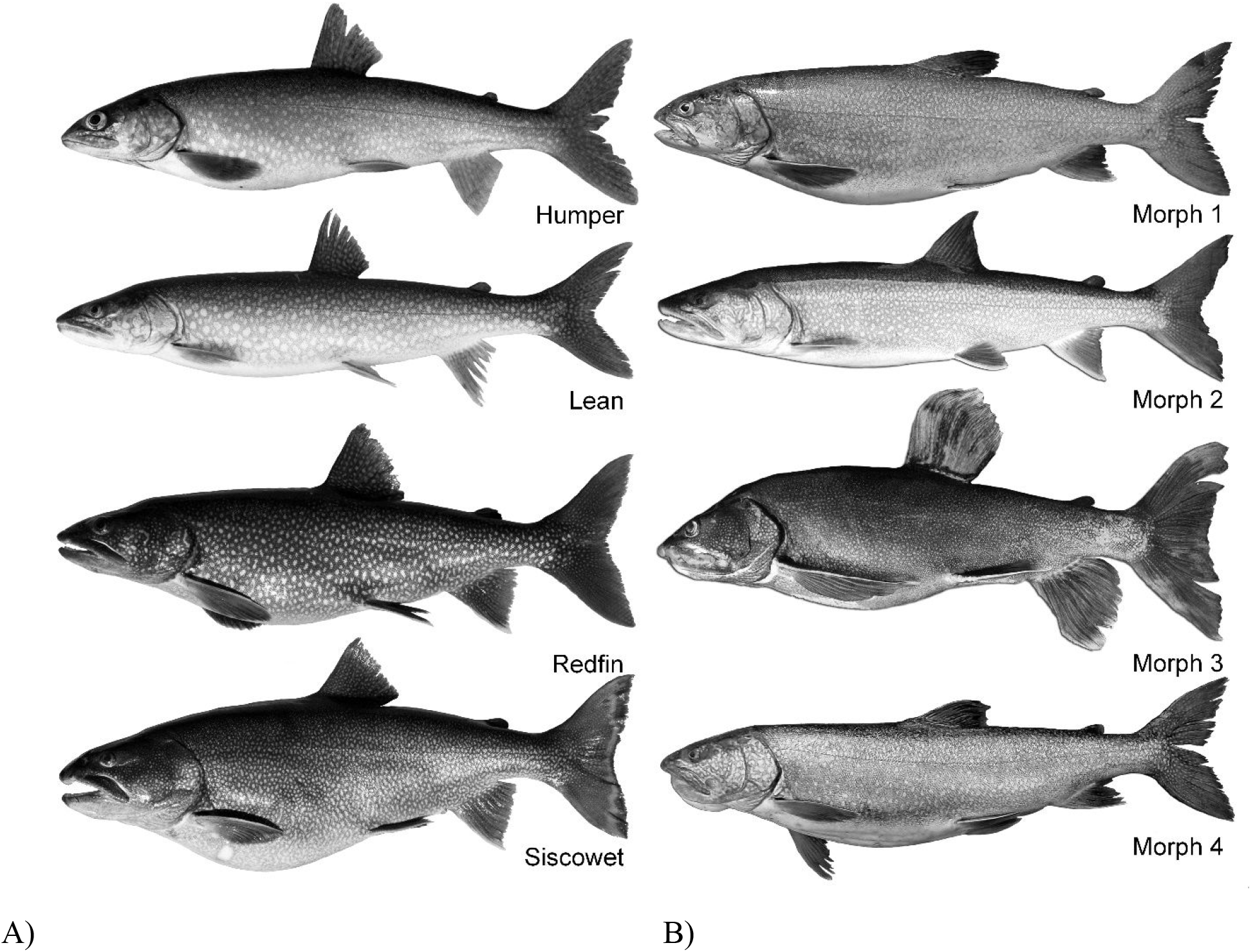
A) The four morphotypes of Lake Trout in Lake Superior, the lean, humper, siscowet, and redfin (defined in Muir et al. 2014). B) The four shallow-water morphotypes of Lake Trout from Great Bear Lake, Morphs1-4 (defined in Chavarie et al. 2013).

Lake Superior is a post-glacial, oligotrophic lake between Canada and the USA (47°43′ N, 86°56′ W) (Fig. 1). Lake Superior is the largest lake in the world by surface area (82 100 km^2^). Most of the waters of Lake Superior can be classified as offshore; 77% of the total area is greater than 80 m deep (maximum depth = 406 m) (see Gorman et al. 2012, Hoffmann 2017, Horns et al. 2003). Lake Superior supports 87 fish species, with lake trout as a main target of the commercial fishery since the 1800s. Lake trout were sampled from two sites, Superior Shoal (48° 3’43.54” N, 87° 8’52.57” W) and Stannard Rock (47°12’26.26” N, 87°12’3.82” W) (Fig. 1).

Lake Superior supports one of the highest levels of sympatric diversity expressed within lake trout (Fig. 2). Four ecotypes are currently recognized; the siscowet, humper, and redfin ecotypes inhabit deep-water (> 70 m), whereas the lean ecotype occupies shallow-water habitats (< 70 m) (Bronte et al. 2003, Bronte and Moore 2007, Muir et al. 2014). The siscowet is characterized by a large head, short snout, long maxilla, large eye, short and deep caudle peduncle, and moderately long paired fins (Muir et al. 2015). Humpers have a small head, short snout, short maxillae, large eyes, and short and narrow caudal peduncle. (Moore and Bronte 2001, Muir et al. 2015). The redfin has the largest head, snout, and eyes, the longest and deepest caudal peduncle, and much longer pelvic and pectoral fins than the other ecotypes (Muir et al. 2014). Finally, lean lake trout have a large, narrow, and pointed head, long snout, small eyes, long and narrow caudal peduncle, short paired fins, and low body lipid content. (Endler 1978, Khan and Qadri 1970, Muir et al. 2015). As vertical migrating visual predators, the three deep-water ecotypes are likely feeding mostly on *Mysis* and deep-water ciscoes (*Coregonus artedi* complex) (Hoffmann 2017, Hrabik et al. 2006, Muir et al. 2014), whereas the lean ecotype is adapted for daytime predation on pelagic fishes in shallow-water habitats (piscivorous feeding strategy; Harvey and Kitchell 2000, Harvey et al. 2003, Janhunen et al. 2009).

### Fatty acids

Dorsal muscle samples (Budge et al. 2011) from lake trout in both lakes were stored at −20 °C (Budge et al. 2006, Chavarie et al. 2016b, Kavanagh et al. 2010, Loseto et al. 2009). Lipids were extracted from 1 g of the homogenate material; after passive overnight extraction (at −20) in 2:1 chloroform:methanol containing 0.01% BHT (v/v/w) (Folch et al. 1957), samples were filtered through Whatman Grade 1 Qualitative filter paper and the filter paper sample was rinsed twice with 2 mL of 2:1 chloroform:methanol. Sample extract was collected in a test tube and 7 mL of 0.88 NaCl solution were added to encourage fatty acids to move into the organic (chloroform) layer. The aqueous layer was discarded, after which the chloroform was dried with sodium sulfate prior to total lipid determination. The extracted lipid was used to prepare fatty acid methyl esters (FAME) by transesterification with Hilditch reagent (0.5 N H_2_SO_4_ in methanol) (Morrison and Smith 1964). Samples were heated for 1 h at 100 °C. Gas chromatographic (GC) analysis was performed on an Agilent Technologies 7890N GC equipped with a 30 m J&W DB-23 column (0.25 mm I.D; 0.15 μm film thickness). The GC was coupled to a Flame Ionization Detector operating at 350 °C. Hydrogen was used as carrier gas flowing at 1.25 mL/min for 14 minutes, increasing to 2.5 mL/min for 5 minutes. The split/splitless injector was heated to 260 °C and run in splitless mode. The oven program was as follows: 60 °C for 0.66 min, increasing by 22.82 °C/min to 165 °C with a 1.97 min hold; increasing by 4.56 °C/min to 174 °C and by 7.61 °C/min to 200 °C with a 6 min hold. Peaks were quantified using Agilent Technologies ChemStation software. Fatty acids standards were obtained from Supelco (37 component FAME mix) and Nuchek (54 component mix GLC-463). FAMEs were identified via retention time and known standard mixtures and are reported as percentages of total fatty acids. Fatty acid standards were obtained from Supelco – Oakville Ontario, Canada (37 component FAME mix) and Nuchek – Elysian Minnesota, USA (54 component mix GLC-463). Standards were run to allow creation of a 4 level calibration curve for each set of samples at the start of each sample set. A standard was repeated every 10 samples thereafter. Every 10 th sample was injected in duplicate. All fatty acids values were converted to a mass percentage of the total array, and were named according the IUPAC nomenclature as X:Y n-z, where X is the number of carbons in the fatty acid, Y is the number of methylene-interrupted double bonds present in the chain, and n-z denotes the position of the last double bond relative to the methyl terminus (Ronconi et al. 2010). All laboratory analyses were conducted at the Freshwater Institute, Fisheries and Oceans Canada, Winnipeg MB.

For both lakes, tissue samples were taken from indivual lake trout previously identified to corresponding ecotypes: 126 samples were collected from Great Bear Lake (Ecotype 1 = 32, Ecotype 2 = 35, Ecotype 3 = 38, and Ecotype 4 = 21, see Chavarie et al. 2016b for more details) and 210 samples were collected from Lake Superior (60 siscowet, 60 humper, 30 redfin, and 60 lean, see Hoffmann 2017 for more details). Overall, fatty acid analysis procedures were divided into two steps, using non-dietary and dietary fatty acid biomarkers (see Appendix 1), following the methods of Iverson et al. (1997, 2004) and Budge et al. (2006) to identify dietary fatty acids biomarkers. Dietary biomarkers were defined as fatty acids ≥ 14 carbons that are generally incorporated into animal tissue from the diet with no or little modification (e.g., rather than come from biosynthesis) (Iverson 2009). Thirty-eight dietary and 24 non-dietary fatty acids biomarkers were found to be shared between the two lakes, and these were selected for further analyses (Table A1 and A2).

### Statistical Analyses

Unless noted otherwise, statistical analyses were conducted using R software version 3.5.3 (R Core Team 2017). Prior to analysis, fatty acid concentrations were logit transformed (log(p/(1−p)) to normalize the data, and then scaled and centered using a z-score transformation (z=xμ/σ). (Clemmensen et al. 2011, Witten and Tibshirani 2011). Principal Component Analysis (PCA) was performed on all dietary and non-dietary fatty acid biomarkers to provide inference about patterns of variation among locations and ecotypes (Chavarie et al. 2016b). PCA summarizes similarities and differences among individuals, based on their fatty acid profiles, independent of ecotype and location (Chavarie et al. 2016b). For PCAs, 12 outlier individuals from Superior Shoal were excluded because PCAs are sensitive to outlier variation (but see Fig. A1 for PCAs with outliers included) (Filzmoser et al. 2009, Kriegel et al. 2008).

To test for differences in fatty acid composition among ecotypes within each lake (dietary and non-dietary), we used Permutational Multivariate Analysis of Variance (PERMANOVA; a non-parametric analog of Multivariate Analysis of Variance), followed by post-hoc comparisons with Bonferroni corrections. PERMANOVAs were performed in PAST 3 (Hammer et al. 2001) using 9999 permutations. A similarity percentage routine (SIMPER) using Bray-Curtis was used to determine which fatty acids (dietary and non-dietary) were primarily responsible for observed differences among ecotypes for each lake (King and Jackson 1999). We also performed linear discriminant analysis on fatty acids (dietary and non-dietary) to delineate differences among ecotypes at each location. A jacknife validation procedure, using 20% of our data as unknown, provided a classification success metric to assess how distinct ecotypes appeared in each fatty acids dataset (dietary vs non-dietary).

## Results

### Combined lakes analyses

The first two axes of the PCAs explained 42.4 % and 41.5%, respectively, of the variation among individuals in their non-dietary and dietary fatty acid biomarkers (Fig. 3). In both PCAs, lake trout from Great Bear Lake were largely distinct from Lake Superior trout (only ~30 individuals from Great Bear Lake overlapped with individuals from Lake Superior), but trout from the two Lake Superior sites, Stannard Rock and Superior Shoal, overlapped. The non-dietary fatty acids 13:1, 14:1n-7, 15:0, 15:1n8, 15:1n6, 15:0 iso, 16:1n11, 17:0 iso, 20:0, 22:0, 20:2n9, and 24:1n9 contributed to the separation between the two lakes. Separation between the two lakes in the dietary PCA appeared to be driven by fatty acids associated with pelagic habitat (14:0, 20:1n-9, 20:1n-7, 20:1n-11, and 22:1n-9; toward Lake Superior) versus one dietary fatty acid associated with cannibalism or/and carnivory (20:5n-3; toward Great Bear Lake) (Appendix, Table 1). Finally, the first two axes of the PCA based on all fatty acids combined explained 39.0% of the variation among lake trout ecotypes from Great Bear Lake and Lake Superior. As before, lake trout from Great Bear Lake were largely separated from the Lake Superior trout, whereas lake trout from Stannard Rock and Superior Shoal in Lake Superior overlapped completely.

**Table 1.**
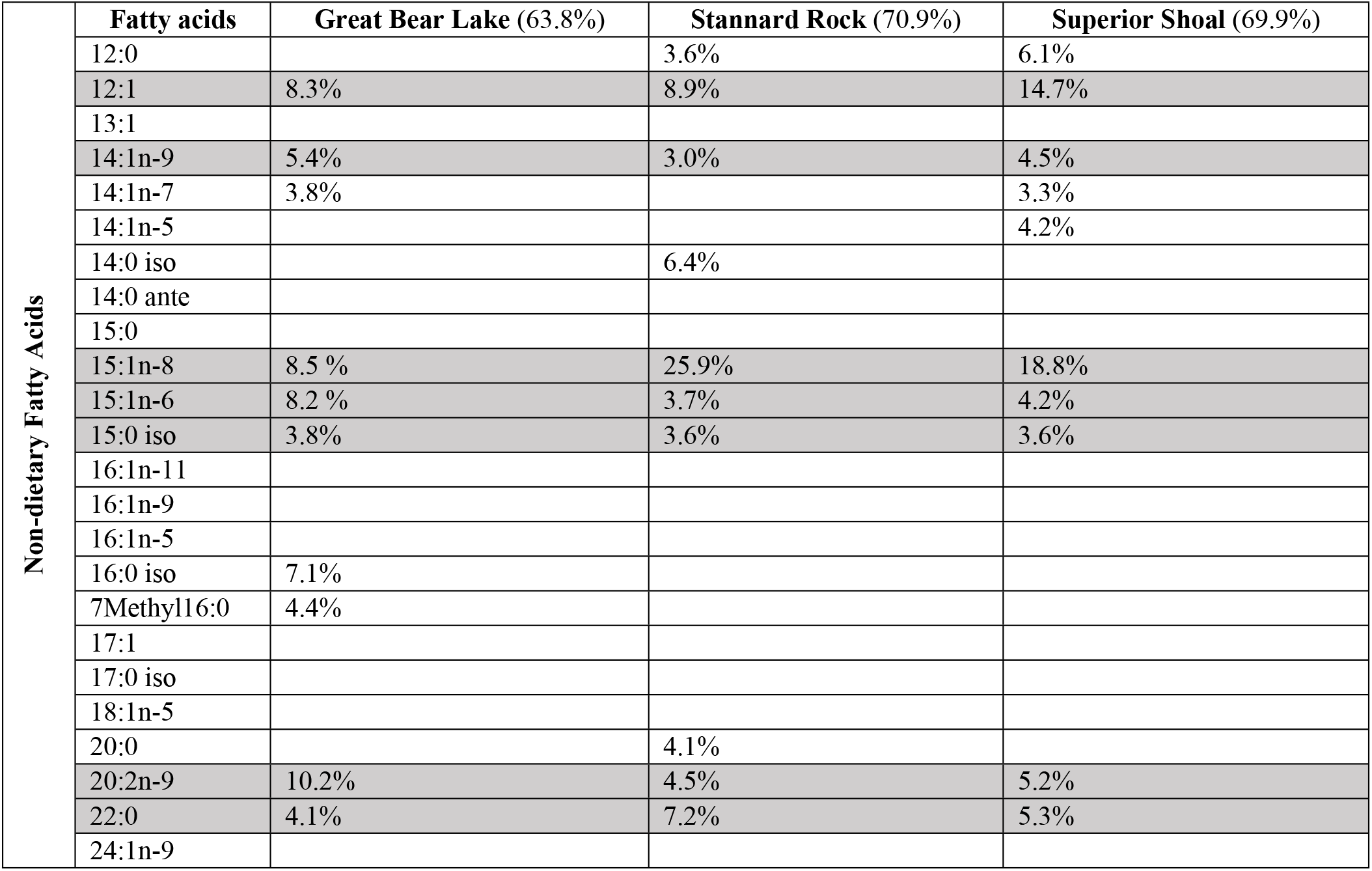
The ten most discriminating non-dietary fatty acids biomarkers from SIMPER analyses to determine which fatty acids were primarily responsible for observed differences. Results are presented for each region, including percentage contribution to overall fatty acid dissimilarity among lake trout morphs. Fatty acids are listed in order of elution; those highlighted in grey are shared among the study three regions. The total percentage of the ten most discriminating fatty acids are given for each region.

| Non-dietary Fatty Acids | Fatty acids | Great Bear Lake (63.8%) | Stannard Rock (70.9%) | Superior Shoal (69.9%) |
| --- | --- | --- | --- | --- |
|  | 12:0 |  | 3.6% | 6.1% |
|  | 12:1 | 8.3% | 8.9% | 14.7% |
|  | 13:1 |  |  |  |
|  | 14:1n-9 | 5.4% | 3.0% | 4.5% |
|  | 14:1n-7 | 3.8% |  | 3.3% |
|  | 14:1n-5 |  |  | 4.2% |
|  | 14:0 iso |  | 6.4% |  |
|  | 14:0 ante |  |  |  |
|  | 15:0 |  |  |  |
|  | 15:1n-8 | 8.5 % | 25.9% | 18.8% |
|  | 15:1n-6 | 8.2 % | 3.7% | 4.2% |
|  | 15:0 iso | 3.8% | 3.6% | 3.6% |
|  | 16:1n-11 |  |  |  |
|  | 16:1n-9 |  |  |  |
|  | 16:1n-5 |  |  |  |
|  | 16:0 iso | 7.1% |  |  |
|  | 7Methyl16:0 | 4.4% |  |  |
|  | 17:1 |  |  |  |
|  | 17:0 iso |  |  |  |
|  | 18:1n-5 |  |  |  |
|  | 20:0 |  | 4.1% |  |
|  | 20:2n-9 | 10.2% | 4.5% | 5.2% |
|  | 22:0 | 4.1% | 7.2% | 5.3% |
|  | 24:1n-9 |  |  |  |

**Fig. 3.**
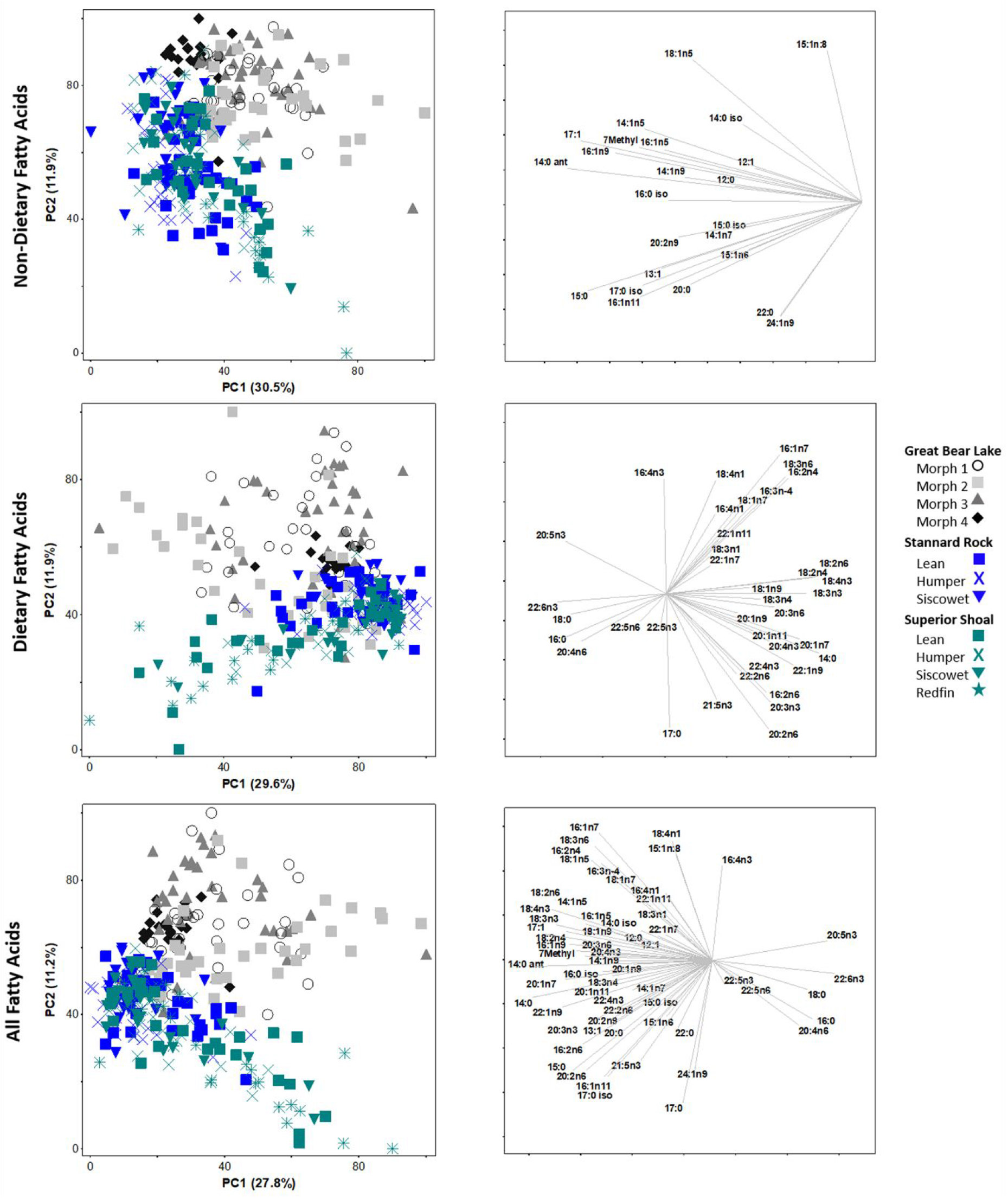
PCA of non-dietary fatty acids biomarkers (top panels), dietary fatty acids biomarkers (middle panels), and all fatty acids (bottom panels) for individual Lake Trout collected from Great Bear Lake, Stannard Rock (Lake Superior), and Superior Shoal (Lake Superior). Vectors of individual fatty acids important to the positioning of lake trout are represented to the right of each PCA. Angles and lengths of vectors represent the direction and strength of relationships, respectively, between variables and the principal components.

### Lake Trout intraspecific diversity of Great Bear Lake

Composition of both non-dietary and dietary fatty acid biomarkers were able to discriminate among the four lake trout ecotypes from Great Bear Lake. Non-dietary fatty acid biomarkers differed among the four Great Bear Lake ecotypes (one-way PERMANOVA, F_3,122_ = 3.6, P < 0.01). Similarly, comparison of non-dietary fatty acids biomarkers showed that all pairs of ecotypes differed from one another (P < 0.01) except for ecotypes 1 and 3. The ten most discriminating non-dietary fatty acid biomarkers from SIMPER explained 63.8% of the dissimilarity among groups (Table 1). The first two axes of the linear discriminant analysis explained 55.7 % and 27.8% of the variation, and 57.9% of all individuals were correctly classified to ecotype based on non-dietary fatty acids biomarkers (Fig. 4). Dietary fatty acid biomarkers also differed among the four ecotypes from Great Bear Lake (one-way PERMANOVA, F_3,122_ = 2.95, P < 0.01), and most ecotypes differed from each other (all pairwise P < 0.01 except for ecotypes 1 vs. 3). The ten most discriminating dietary fatty acid biomarkers from SIMPER explained 48.8% of the dissimilarity among groups (Table 2). The first two axes of the linear discriminant analysis explained 46.7% and 39.0% of the variation, and 68.3% of all individuals were correctly classified to ecotype based on dietary fatty acids biomarkers (Fig. 4).

**Table 2.** The ten most discriminating dietary fatty acids biomarkers from SIMPER analyses to determine which fatty acids were primarily responsible for observed differences. Results are presented for each region, including percentage contribution to overall fatty acid dissimilarity among lake trout morphs. Fatty acids are listed in order of elution; those highlighted in grey are shared among the study three regions. The total percentage of the ten most discriminating fatty acids are given for each regions.

| Dietary and Extended-Dietary Fatty Acids | Fatty acids | Great Bear Lake (48.8%) | Stannard Rock (55.2%) | Superior Shoal (55.8%) |
| --- | --- | --- | --- | --- |
|  | 14:0 |  |  |  |
|  | 16:0 |  |  |  |
|  | 16:1n-7 |  |  |  |
|  | 16:2n-6 |  |  |  |
|  | 16:2n-4 |  |  |  |
|  | 17:0 |  |  |  |
|  | 16:3n-4 | 4.0% | 2.5 % | 2.9% |
|  | 16:4n-3 | 3.0% | 2.6% |  |
|  | 16:4n-1 |  | 7.3% | 6.3% |
|  | 18:0 |  |  |  |
|  | 18:1n-9 |  | 2.4% |  |
|  | 18:1n-7 |  |  |  |
|  | 18:2n-6 |  |  |  |
|  | 18:2n-4 |  |  |  |
|  | 18:3n-6 |  |  |  |
|  | 18:3n-4 |  |  |  |
|  | 18:3n-3 |  |  |  |
|  | 18:3n-1 | 3.3% |  | 3.5% |
|  | 18:4n-3 |  |  |  |
|  | 18:4n-1 | 3.5% | 10.5% | 11.1% |
|  | 20:1n-11 | 4.0% |  |  |
|  | 20:1n-9 |  |  |  |
|  | 20:1n-7 | 3.6% |  |  |
|  | 20:2n-6 |  |  |  |
|  | 20:3n-6 |  |  |  |
|  | 20:4n-6 |  |  |  |
|  | 20:3n-3 |  |  |  |
|  | 20:4n-3 |  |  |  |
|  | 20:5n-3 |  |  |  |
|  | 22:1n-11 | 8.6% | 11.9% | 12.1 % |
|  | 22:1n-9 |  |  |  |
|  | 22:1n-7 |  | 2.7% | 4.6% |
|  | 22:2n-6 | 6.3% |  |  |
|  | 21:5n-3 | 7.5% |  | 4.7 % |
|  | 22:5n-6 |  | 7.2% | 3.0% |
|  | 22:4n-3 | 5.0% | 2.6% | 4.2% |
|  | 22:5n-3 |  |  |  |
|  | 22:6n-3 |  | 5.5% | 3.4% |

**Fig. 4.**
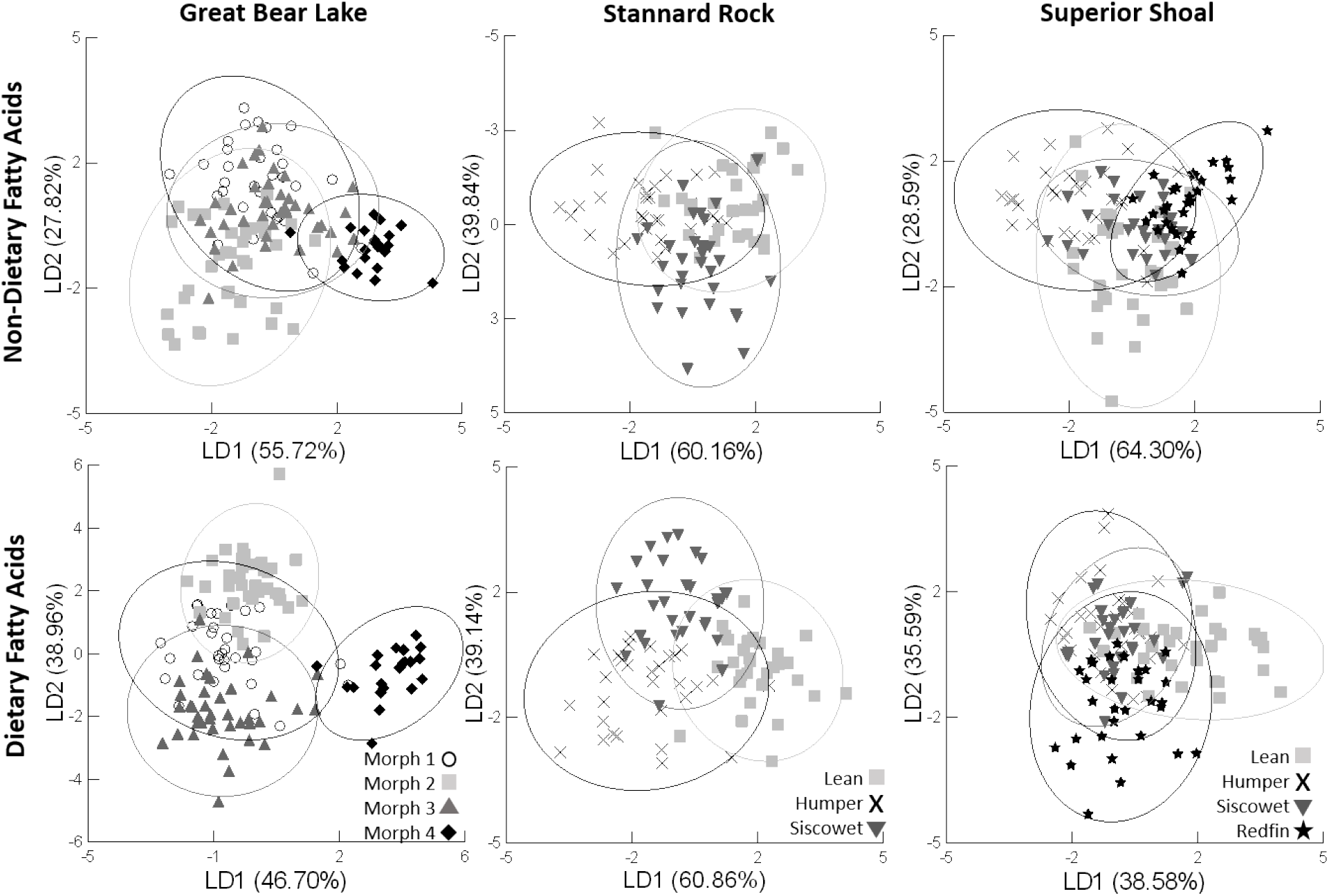
Results of linear discriminant function analyses of non-dietary fatty acids biomarkers (top panel) and dietary fatty acids biomarkers (bottom panel) for Lake Trout collected from Great Bear Lake, Stannard Rock (Lake Superior), and Superior Shoal (Lake Superior). The 95% ellipse of each morph is also provided.

### Lake Trout intraspecific diversity of Lake Superior, Stannard Rock

Non-dietary fatty acid biomarkers differed among the three ecotypes of lake trout from Stannard Rock (one-way PERMANOVA, F_2,87_ = 3.9, P < 0.01), and leans differed significantly from both siscowets and humpers (P < 0.01). The ten most discriminating non-dietary fatty acids biomarkers from SIMPER explained 70.9% of the dissimilarity among ecotypes (Table 1). The first two axes of the linear discriminant analysis explained 60.2% and 39.8% of the variation, respectively, and 52.2% of all individuals were correctly classified to ecotype based on non-dietary fatty acids biomarkers (Fig. 4). In contrast to results from non-dietary fatty acid biomarkers, no differences in dietary fatty acid composition occurred among the three lake trout ecotypes from Stannard Rock, Lake Superior (one-way PERMANOVA, F_2,87_ = 1.12, P = 0.3). The ten most discriminating fatty acids from SIMPER explained 55.2 % of the dissimilarity among groups (Table 2), and the first two axes of the linear discriminant analysis explained 60.9% and 39.1% of the variation. Fifty percent of individuals were correctly classified to ecotype based on dietary fatty acids biomarkers (Fig. 4).

### Lake Trout intraspecific diversity of Lake Superior, Superior Shoal

Like Stannard Rock, composition of non-dietary fatty acid biomarkers differed significantly among the four ecotypes (one-way PERMANOVA, F_3,116_ = 2.5, P < 0.01); leans differed from redfins (P < 0.01) and differences were marginally between redfin and siscowet and between redfin and humper (P < 0.06). The ten most discriminating non-dietary fatty acids biomarkers from SIMPER explained ~69.9% of the dissimilarity among groups (Table 1). The first two axes of the linear discriminant analysis explained 64.3% and 25.6% of the variation, and 45.0% individuals were correctly classified to ecotype using non-dietary fatty acids biomarkers (Fig. 4). Similar to what was found at Stannard Rock, we found no differences among the four ecotypes from Superior Shoal based on dietary fatty acid composition (one-way PERMANOVA, F_3,116_ = 0.8, P = 0.3). The ten most discriminating fatty acids from SIMPER explained 55.8 % of the dissimilarity among groups (Table 2). The first two axes of the linear discriminant analysis explained 38.6% and 35.6% of the variation, respectively, and 31.7% of fish were correctly classified to ecotype based on their dietary fatty acids (Fig. 4).

## Discussion

In this study, non-dietary fatty acids showed variation geographically (between lakes), between locations within a lake (i.e., Stannard Rock versus Superior Shoal), and among ecotypes within a lake or location. Although some overlap existed (which reduced the power to discriminate), our results showed that when investigating intraspecific diversity, non-dietary fatty acids biomarkers can be a useful tool to delineate groups, and that sometimes, such as at sites in Lake Superior, were more discriminatory than dietary fatty acids. While characterizing trophic divergence, food-web linkages, and time-integrated niche use across a large array of taxa, most investigators use a common set of fatty acids that are selected from lists that classify fatty acids as “dietary” or “extended dietary” biomarkers (Budge et al. 2006, Iverson et al. 2004). However, our results suggest that discarding non-dietary fatty acid data as a matter of course may result in inadvertent loss of information.

Not all fatty acids provide equivalent information about diet due to metabolism and *de novo* synthesis (Iverson et al. 1993, Iverson et al. 2004). Metabolism, however, can differ within species, as physiological differences often exists among sets of individuals, i.e., “ecotypes” (Miles et al. 2007, Pryke et al. 2007). Such physiological differences within a species may result in non-dietary fatty acids biomarkers being a relevant tool to delineate intraspecific diversity, as we show here for lake trout. In Lake Superior, fat content has long been recognized as an important characteristic that distinguishes lake trout ecotypes along a depth axis (Eschmeyer and Phillips 1965, Eshenroder 2008), and these lipid differences have been linked to genetic differences among ecotypes (Eschmeyer and Phillips 1965, Goetz et al. 2013, Goetz et al. 2010). Thus, fatty acid deposition and metabolism appear to have undergone selection along a depth gradient likely in part to contribute to buoyancy compensation, with differences reported between siscowet (deep-water ecotype) and lean (shallow-water ecotype) lake trout (Eschmeyer and Phillips 1965, Goetz et al. 2010). Until now, no information has been available for redfin and humper ecotypes.

In addition to the variation in lipid accumulation, Goetz et al. (2013) also found differences in lipid composition between siscowet and lean ecotypes. Because a common garden design was used in their experiment, the higher proportion of polyunsaturated fatty acids (PUFAs) found in the muscle lipid profile of siscowet than in lean lake trout could not be attributed to differences in diet. The differences in non-dietary fatty acids biomarkers we observed among the four ecotypes of lake trout from Lake Superior is consistent with the concept of metabolotypes (Goetz et al. 2013). Altogether, differences in energy processing and storage between lean and siscowet lake trout in Lake Superior are adaptive to their respective habitats, deep-vs. shallow-water, and to their life-histories (Goetz et al. 2013). Lipid content, intertwined with buoyancy variations, have been linked to differences in depth distributions and swimming tactics among lake trout ecotypes (Zimmerman et al. 2006, Zimmerman et al. 2007, Zimmerman et al. 2009). While deep-water ecotypes use hydrostatic lift related to lipid content to enhance vertical migration while foraging for *Mysis* and cisco (Henderson and Anderson 2002), the shallow-water lean ecotype likely relies more on hydrodynamic lift, linked to the cruising movements of pelagic predators (Webb 1984). Consistent with the findings of Goetz et al. (2013), we found that non-dietary fatty acid biomarkers differed between shallow- and deep-water ecotypes at Stannard Rock. However, at Superior Shoal, non-dietary fatty acids biomarkers differed only between the shallow-water lean ecotype and one deep-water ecotype – the redfin. This observation, along with some more subtle differences in non-dietary fatty acid biomarkers among deep-water ecotypes, requires further study as our current knowledge is limited. It is presently unclear if these lake trout ecotypes have adapted physiologically to their habitat leading to metabolotypes and how diets differ among all ecotypes temporally and spatially within Lake Superior. As such, feeding on the same item by different ecotypes might produce different fatty acid accumulations resulting not only in a physiologically interesting questions but also questioning the accuracy of predictions about diet composition.

The concept that dynamics of energy processing and storage are adaptive along a gradient associated with depth does not apply to the intraspecific diversity of lake trout in Great Bear Lake. This diversity is limited to shallow-water habitat, and appears to be independent of major habitat and resource partitioning (Chavarie et al. 2016a, Chavarie et al. 2018). The similarity of results between Great Bear Lake and Lake Superior is thus perplexing, as we were expecting greater differences in non-dietary fatty acid biomarkers among ecotypes in Lake Superior than in Great Bear Lake, due to known buoyancy variation associated with a depth gradient in Lake Superior. If lake trout ecotypes from Great Bear Lake are also under selection (Harris et al. 2014), differences in energy processing and storage may be as pronounced as those that have been observed in Lake Superior. Another question raised by our results is the extent to which ecotypes are independent of major habitat or resource axes (the same question is pertinent for Lake Superior), especially because dietary fatty acid biomarkers in Great Bear Lake were slightly better at delineating intraspecific groups than non-dietary biomarkers than in Lake Superior.

Differences between morphs and sites may be influenced by the total content of fatty acids (μg FA/m) in muscle, rather than by their composition (%). Tissue-specific storage of fatty acids can vary across space or time, and some fish store lipids as modified adipose or lipid pockets in their muscle (e.g., salmonids; Iverson 2009, Sasaki et al. 1989). In a comparison of fatty acid content between dorsal muscle vs. belly tissue, (Happel et al. 2019) found a threshold response; the tissues became increasingly dissimilar when lipid content of muscle was > ~10%. Nevertheless, Happel et al. (2019) found that fatty acid profiles were specific to each of the five lakes they examined, i.e., lake trout displayed broad variation among locations.

Ecological differences in allopatry are often more pronounced than those in sympatry, due to disparate environments and isolation (Fraser et al. 2011, Heggenes 2002, Rundle and Nosil 2005, Yoder et al. 2010). Our results were consistent with this trend, i.e., differences between lakes were greater than differences among ecotypes within a lake for both dietary and non-dietary fatty acid biomarkers. For dietary fatty acids, differences between the two lakes appeared to be due to fatty acids associated with a pelagic environment, such as C20 and C22 monounsaturates, that can be used as biomarkers of food webs based on pelagic copepods (Ahlgren et al. 2009, Budge et al. 2006, Dalsgaard et al. 2003). Although benthic productivity generally dominates in Arctic lakes (Chavarie et al. 2018, Johnson 1975), few lake trout in Great Bear Lake specialized on pelagic resources (i.e., few ecotypes 2 and 4 had fatty acid signatures that overlapped with lake trout from Lake Superior; Chavarie et al., submitted; Chavarie et al. 2016a, Chavarie et al. 2016b).

With the general benthic orientation of lake productivity in Arctic regions, combined with the known distributions in deep-water habitats of lake trout ecotypes from Lake Superior (50-150 m; Muir et al. 2014), our overall results reflected the expected ecological differences of this species from these two systems. Despite these inter-lake differences, similar dietary (e.g., five out of 10) and non-dietary (e.g., 7 out of 10) fatty acid biomarkers were important in identifying intraspecific diversity within each lakes. The greater number of shared non-dietary fatty acids biomarkers discriminating lake trout intraspecific diversity in the two lakes supports the idea of similar physiological differences (e.g., energy processing and storage) among ecotypes from both lakes.

A few caveats should be noted that could alter interpretations of this study. First, we cannot ensure the fatty acid biomarkers defined based on Iverson et al. (2004) truly reflect dietary vs non-dietary origins, due to the lack of taxa-specific studies on the integration of prey fatty acids. Despite this uncertainty, the aim of this paper was to examine loss of information from discarding fatty acids generally considered as non-dietary biomarkers by the literature. Second, no spatial component was defined for the Great Bear Lake dataset, due to small sample sizes from each location, which may have introduced variation into the results. The importance of spatial variation in large, complex systems was shown here for Lake Superior (Chavarie et al. 2015, Hoffmann 2017) and environmental variables (e.g., temperature, light), which may vary spatially, are known to alter lipid composition in fish tissue (Olsen 1999). Third, multiple sizes and life-stages, e.g., juvenile, mature, and resting individuals, were included in the Lake Superior analysis; different life-stages can vary in lipid metabolism (Sheridan 1989) and large lake trout can rely more on nearshore-benthic food web resources than small lake trout (Happel et al. 2017a). Finally, some fatty acids exist at very low amounts (≤ 2%), which can introduce error when interpreting differences among fatty acids that are found in only trace amounts (e.g., peak shouldering) (Christie 1998). Despite these limitations, we found some consistent patterns with regards to intraspecific diversity between lakes and among ecotypes within lakes from two distinct datasets.

## Conclusion

Our study demonstrated the potential benefits of using both dietary and non-dietary fatty acid biomarkers for delineating variation within a species. In some instances, non-dietary fatty acids were better for discriminating ecotypes than dietary ones, in contrast to the popular maxim associated with trophic markers, “you are what you eat”. Dietary fatty acid biomarkers can document the occurrence of discrete niche use among sets of individuals (i.e., ecotypes) within a species (Chavarie et al. 2016b). The fatty-acid composition of individuals that reflects their diet has been validated for lake trout (Happel et al. 2016a, Happel et al. 2016b). However, physiological differences in the dynamics of energy processing (e.g., metabolism) and storage can also be adaptive among ecotypes (Eschmeyer and Phillips 1965, Goetz et al. 2013, Goetz et al. 2010), which may result differences in non-dietary fatty acids (e.g., greater than dietary fatty acids) and thus, be useful molecular tools. The lack of information on non-dietary fatty acids in lake trout (and other taxa), however, raises questions about their physiological role and if consumers acquire them or not from their diet (e.g., synthesis de novo, elongation). This would be a fruitful area for future research.

Ultimately, the relative importance of dietary vs. non-dietary fatty acid biomarkers depends on the question being asked. In our study, non-dietary fatty acids were valuable in delineating intraspecific variation within a lake, but also in examining differences between lakes. Non-dietary fatty acid biomarkers can provide useful information; therefore, one should carefully consider if such information is superfluous or not before data from these fatty acids are discarded.

## Acknowledgement

We thank Déline Renewable Resources Council, Déline Lands and Finance Corporation, the community of Déline, DFO in Hay River, and the Department of Environment and Natural Resources in Déline, which provided valuable help with field planning and logistics. Financial support was provided by Fisheries and Oceans Canada (DFO), Natural Sciences and Engineering Research Council of Canada, Sahtu Renewable Resource Board, Association of Canadian Universities for Northern Studies, Canadian Circumpolar Institute’s Circumpolar/Boreal Alberta Research and Northern Scientific Training Program, D. Alan Birdsall Memorial Scholarship Fund, Aboriginal Affairs and Northern Development Canada Northwest Territories Cumulative Impacts Monitoring Program grants, and the Great Lakes Fishery Commission. Logistical and in-kind support were provided by the Polar Continental Shelf Program and USGS. The findings and conclusions in this article are those of the authors and do not necessarily represent those of the U.S. Geological Survey or the U.S. Fish and Wildlife Service.

## Data Availability Statement

All data presented are available by request via e-mail to the first author.

## Conflict Of Interest

The authors declare that they have no conflict of interest.

## Appendix

**Table A1.** List of 38 of 41 fatty acids shared by the two datasets, considered as either “dietary” fatty acids or “extended-dietary” fatty acids biomarkers and used in this study (see Iverson *et al*., 2004), and the dietary component they are associated with, based on literature: Sargent *et al*., 1995; Brett & Müller-Navarra, 1997; Kattner *et al*., 1998; Virtue *et al*., 2000; Budge et al., 2002; Dalsgaard *et al*., 2003; Iverson *et al*., 2004; Käkelä *et al*., 2005; Alfaro *et al*., 2006; Tucker *et al*., 2008; Ahlgren *et al*., 2009; Gladyshev *et al*., 2009; Loseto *et al*., 2009; Stowasser *et al*., 2009; Piché *et al*., 2010; Mariash *et al*., 2011. The fatty acids highlighted are the one discarded because they were not quantified for both lakes.

| Dietary Fatty acids | Dietary component |
| --- | --- |
| 14:0 | Pelagic (Zooplankton) + diatom |
| 16:0 | Cannibalism or/and carnivorous |
| 16:1n7 | Benthic (bacterial synthesis + diatoms)<br>Cannibalism or/and carnivorous |
| 16:2n6 |  |
| 16:2n4 | Diatom |
| 17:0 |  |
| 16:3n-4 |  |
| 16:4n3 |  |
| 16:4n1 | Diatom |
| 18:0 | Cannibalism or/and carnivorous |
| 18:1n9 | Pelagic (Zooplankton) |
| 18:1n7 | Benthic (bacterial synthesis + algal) |
| 18:2n6 | Terrestrial |
| 18:2n4 |  |
| 18:3n6 |  |
| 18:3n4 |  |
| 18:3n3 | Terrestrial |
| 18:3n1 |  |
| 18:4n3 | Pelagic (Zooplankton) |
| 18:4n1 |  |
| 20:0 | *not used |
| 20:1n11 | Copepod (Iverson, 2009) |
| 20:1n9 | Pelagic (Calanoid copepods diet based) |
| 20:1n7 | Pelagic (Zooplankton) |
| 20:2n9 | *not used |
| 20:2n6 |  |
| 20:3n6 |  |
| 20:4n6 | Benthic (diatom) |
| 20:3n3 |  |
| 20:4n3 |  |
| 20:5n3 | Cannibalism or/and carnivorous |
| 22:1n11 | Copepod |
| 22:1n9 | Pelagic (Zooplankton) |
| 22:1n7 |  |
| 22:2n6 |  |
| 21:5n3 |  |
| 22:4n6 | *not used |
| 22:5n6 | Pelagic (Calanoid copepods diet based) |
| 22:4n3 |  |
| 22:5n3 |  |
| 22:6n3 | Benthic (Pennate diatoms + dinoflagellates +<br>bivalves)<br>Cannibalism |

**Table A2.** List of 24 fatty acids considered as “non-dietary” biomarkers in this study.

| Non-Dietary Fatty acids |
| --- |
| 12:0 |
| 12:1 |
| 13:1 |
| 14:1n9 |
| 14:1n7 |
| 14:1n5 |
| 14:0 iso |
| 14:0 ante |
| 15:0 |
| 15:1n8 |
| 15:1n6 |
| 15:0 iso |
| 16:1n11 |
| 16:1n9 |
| 16:1n5 |
| 16:0 iso |
| 7Methyl 16:0 |
| 17:1 |
| 17:0 iso |
| 18:1n5 |
| 20:0 |
| 20:2n9 |
| 22:0 |
| 24:1n9 |

**Fig. A1.**
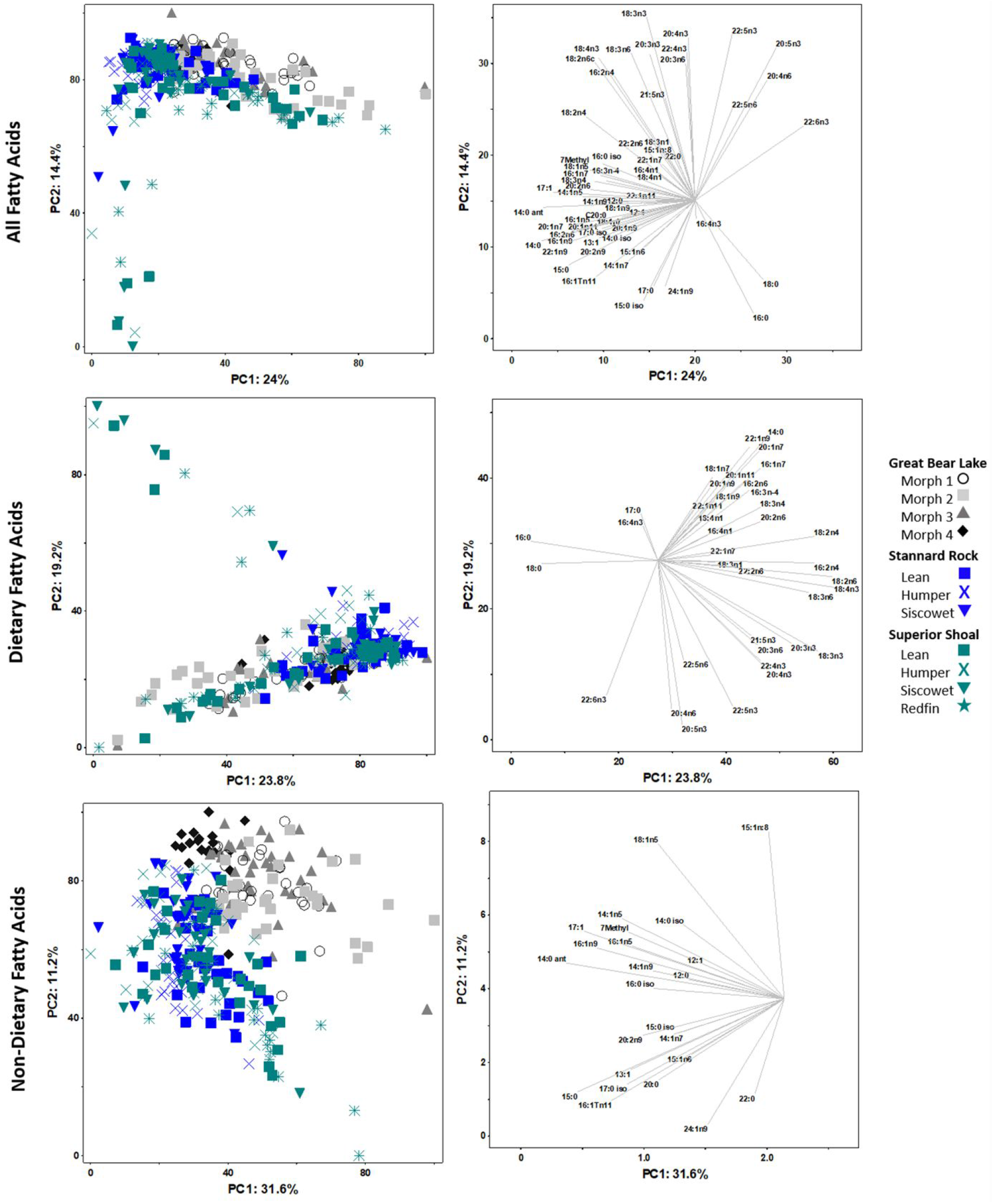
PCA of all fatty acids (top panels), dietary fatty acids biomarkers (middle panels), and non-dietary fatty acids biomarkers (bottom panels) of each morph of Lake Trout from Great Bear Lake, Stannard Rock (Lake Superior), and Superior Shoal (Lake Superior), including the 12 outlier individuals. Vectors of individual fatty acids important to the positioning of lake trout are represented to the right of each PCA. Angles and lengths of vectors represent the direction and the strength, respectively, of relationships between variables and the principal components.

## Notes

https://www.nrcresearchpress.com/doi/10.1139/cjfas-2019-0343#.XosArIhKiUm

